# Different preference functions act in unison: mate choice and risk-taking behaviour in the Atlantic molly (*Poecilia mexicana*)

**DOI:** 10.1101/790832

**Authors:** Carolin Sommer-Trembo, Michael Schreier, Martin Plath

## Abstract

Consistent individual differences in behaviour (animal personality) are widespread throughout the Animal Kingdom. This includes variation in risk-taking versus risk-averse behavioural tendencies. Variation in several personality dimensions is associated with distinct fitness consequences and thus, may become a target of natural and/or sexual selection. However, the link between animal personality and mate choice—as a major component of sexual selection—remains understudied. We asked (1) whether females and males of the livebearing fish *Poecilia mexicana* prefer risk-taking mating partners (directional mating preference), (2) or if their preferences are dependent on the choosing individual’s own personality type (assortative mating). We characterized each test subject for its risk-taking behaviour, assessed as the time to emerge from shelter and enter an unknown area. In dichotomous association preference tests, we offered two potential mating partners that differed in risk-taking behaviour but were matched for other phenotypic traits (body size, shape, and colouration). Females, but not males, exhibited a strong directional preference for risk-taking over risk-averse mating partners. At the same time, the strength of females’ preferences correlated positively with their own risk-taking scores. Our study is the first to demonstrate that a strong overall preference for risk-taking mating partners does not preclude effects of choosing individuals’ own personality type on (subtle) individual variation in mating preferences. More generally, two different preferences functions appear to interact to determine the outcome of individual mate choice decisions.

## Introduction

Behavioural differences among individuals that are consistent over time and across contexts (animal personality or temperament) can be found throughout the Animal Kingdom (Gosling, 2001; Kralj-Fišer & Schuett, 2014; Weiss, 2018). Additive genetic effects contribute to variation in animal personality traits (Dochtermann, Schwab & Sih, 2015), and heritability estimates are often comparable to those reported for life-history and physiological traits (Dochtermann, Schwab, Anderson Berdal, Dalos & Royauté, 2019). This renders animal personality a potential target of both natural and sexual selection (Dingemanse & Réale, 2005; Schuett, Tregenza & Dall, 2010). Accordingly, several empirical studies demonstrated fitness consequences for different personality dimensions, such as boldness (risk-taking behaviour), exploration and sociability (Ballew, Mittelbach & Scribner, 2017; Cote, Dreiss & Clobert, 2008; Dingemanse, Both, Drent & Tinbergen, 2004).

A meta-analysis across taxonomic groups suggested that individual variation along the continuum between risk-taking and risk-averse behaviour—the most studied animal personality dimension—tends to be associated with an increased reproductive success of risk-taking individuals, particularly males (Smith & Blumstein, 2008). It remains controversial, however, what mechanisms might explain fitness variation among behavioural types and whether sexual selection (e.g. female choice in favour of risk-taking mating partners) plays a role in this context (Schuett, Tregenza & Dall, 2010; Smith & Blumstein, 2008). Female guppies (*Poecilia reticulata*), for example, preferred risk-taking over risk-averse males when risk-taking tendencies were experimentally manipulated by presenting single males close to (or away from) predators (Godin & Dugatkin, 1996). The question remains as to whether females would be able to assess actual male personality types independent of males’ behavioural responses to predators. Females could base their mate choice on correlated phenotypic traits, as risk-taking behavioural tendencies (and other personality traits) can be associated with variation in traits like body colouration, or size (Brown, Jones & Braithwaite, 2007; Schweitzer, Montreuil & Dechaume-Moncharmont, 2015). However, even if those phenotypic traits are carefully matched between experimentally presented potential mating partners—as we did in our present study (see below)—systematic co-variation between risk-taking behavioural tendencies and other behaviours could be used for mate assessment. This includes differences in swimming patterns (Kern, Robinson, Gass, Godwin & Langerhans, 2016; Wilson & Godin, 2009), body posture, or readiness to resume normal swimming behaviour after disturbance (Brown, Jones & Braithwaite, 2005; Sommer-Trembo & Plath, 2018; this study).

While some studies suggest the existence of a directional preference for risk-taking mating partners, others reported contrasting patterns in that risk-taking females preferred risk-taking males and *vice versa*, leading to assortative mating (Jiang, Bolnick & Kirkpatrick, 2013). Assortative mating can affect individuals’ reproductive success (Both, Dingemanse, Drent & Tinbergen, 2005; Kralj-Fišer, Sanguino Mostajo, Preik, Pekár & Schneider, 2013; Scherer, Kuhnhardt & Schuett, 2017). For instance, guppy females that were paired with males showing similar risk-taking tendencies had a higher parturition success than females that were paired disassortatively (Ariyomo & Watt, 2013).

Here, we present a test for directional mate choice and/or assortative mating based on individuals’ risk-taking behaviour in the livebearing fish *Poecilia mexicana*. For the first time, we assessed both male and female mating preferences. While the importance of male mate choice is increasingly acknowledged (Edward & Chapman, 2011), studies on male mate choice for female personality types are virtually absent. We used emergence tests (Brown, Jones & Braithwaite, 2005; Sommer-Trembo & Plath, 2018) to assess individuals’ risk-taking behaviour. *Poecilia mexicana* (including the population studied here) has repeatedly been characterized for risk-taking behaviour, and previous studies reported high behavioural repeatability, with R-values ranging between 0.53 and 0.64 (*freezing time after a simulated predator attack*, R = 0.64, Sommer-Trembo et al., 2016a; repeatability across *time to emerge from shelter* and *freezing time after a simulated predator attack*, R = 0.53, Sommer-Trembo & Plath, 2018). Slightly lower, yet significant R-values were reported for the related guppy (*time to emerge from shelter*, R = 0.51, Brown & Irving, 2013; *time to emerge from shelter*, R = 0.51 for females and R = 0.36 for males, Irving & Brown, 2013; *time to emerge from shelter*, R = 0.33, White, Kells & Wilson, 2016) and other poeciliid fishes (e.g., *Gambusia affinis, time to emerge from shelter*, R = 0.29 in Cote, Fogarty, Weinersmith, Brodin & Sih, 2010 and R = 0.39 in Gomes-Silva, Liu, Chen, Plath & Sommer-Trembo, 2017; *Poecilia vivipara, time to emerge from shelter*, R = 0.70, Sommer-Trembo et al. 2016b). For our present study, we initially screened a large number of potential stimulus and focal individuals so as to be able to select stimulus pairs with contrasting behavioural type (see methods). This time-consuming approach led us to decide to not assess behavioural repeatability, but we argue that behavioural repeatability of risk-taking tendencies is well established in our study species.

We performed dichotomous mate choice tests in which focal individuals could choose between two stimulus individuals of the opposite sex that differed in risk-taking tendencies but were matched for other phenotypic traits known to be involved in mate assessment (body size, shape and colouration; Rios-Cardenas & Morris, 2011). This left only behavioural characteristics correlated with risk-taking tendencies as a potential source for mate assessment. We asked whether focal individuals prefer risk-taking over risk-averse mating partners (directional preference) and/or whether a pattern indicative of assortative mating would be uncovered. Either result would indicate that focal individuals were able to assess the behavioural type of potential mating partners within the short time period of our mate choice tests (focal and stimulus individuals were unfamiliar prior to the tests) and without an opportunity to observe interactions with predators.

While predictions may seem to be mutually exclusive when considering the potential occurrence of directional mating preferences or assortative mating, we argue that this is actually not the case: focal individuals [at least females (Godin & Dugatkin, 1996)] could show an overall (directional) preference for risk-taking mating partners. Still, ‘hidden’ within the individual variation in mating preferences, focal individuals’ own risk-taking tendencies might predict the strength at which individuals express this mating preference. Our present study confirms that both preference functions indeed act in unison and jointly explain female (but not male) mate choice for risk-taking mating partners.

## Materials and methods

### Test subjects and general testing procedure

Test subjects were laboratory-reared descendants of wild-caught Atlantic mollies (*Poecilia mexicana*), which we collected in the southern Mexican Río Oxolotán in 2013. We maintained the fish in several aerated and filtered 200-L stock tanks at 28°C under a 12/12 h light/dark cycle. Our stock tanks comprised juveniles and adults of both sexes at densities of 50–70 adult individuals per tank. We fed the fish twice a day *ad libitum*-amounts of commercially available flake food (Tetra Min^®^), frozen spinach, *Artemia* naupliae and frozen bloodworms (*Chironomus* larvae). Aquaria were equipped with live and artificial plants and stones. To maintain water quality, we replaced half of the water by aged tap water every 2 weeks. Focal and stimulus fish for the mate choice tests were taken from different stock tanks and were thus unfamiliar prior to the tests.

We conducted our behavioural experiments in 2016. Before the behavioural assessments, test subjects were held for three days in same-sex groups at densities of 20 individuals per tank. We initially tested a large number of fish (*n* = 300) for risk-taking tendencies, after which they were given three days for recovery before focal individuals and stimulus pairs were selected. We tested a sub-set of *n* = 54 focal individuals (27 females and 27 males) for their mating preferences by using dichotomous mate choice tests (see below for details on which individuals were selected for the mate choice tests).

### Assessment of risk-taking tendencies

We used time to emerge from shelter and enter an unknow area as a proxy of individuals’ risk-taking tendency (Brown, Jones & Braithwaite, 2005; Sommer-Trembo & Plath, 2018). To this end, the test subject was gently transferred into the shelter compartment of the test tank, which was equipped with stones and artificial plants (see Sommer-Trembo & Plath, 2018 for details). After a 3-min acclimatization period (after which all tested individuals showed normal swimming behaviour), we lifted an opaque Plexiglas divider and measured the time until the fish entered the open field area with a uniformly grey bottom and no opportunities for hiding. Based on a pilot study, fish were given a maximum of 60 s to emerge from shelter. We calculated individual risk-taking scores as: [maximum emergence time (60 s) – individual emergence time], which resulted in high sores for risk-taking and low scores for risk-averse individuals. 85% of test subject emerged from shelter within 60 s, whereas the other 15% reached the ceiling value of 60 s.

### Assignment of focal and stimulus individuals

Initially, we assessed risk-taking tendencies of *n* = 150 females and *n* = 150 males. We randomly selected 60 of those individuals (*n* = 30 per sex), based on the flip of a coin, which did not undergo any further behavioural test before they served as focal individuals (see below). Of the remaining 240 individuals, we disregarded individuals with intermediate boldness-scores (emergence times between 21 and 39 s) and retained those individuals as potential stimulus fish that could be characterized unambiguously as either risk-taking (emergence times ≤ 20 s) or risk-averse (≥ 40 s).

We measured the standard length (SL) of all individuals meeting these criteria to the nearest millimetre by laying the fish flat on moist laminated millimetre paper and matched stimulus pairs according to their SL (difference ≤ 3 mm). Body size is known to be an important criterion of mate choice in poeciliids (Bisazza, Marconato & Marin, 1989; Herdman, Kelley & Godin, 2004; Plath, Seggel, Burmeister, Heubel & Schlupp, 2006). Additionally, we visually matched the respective stimulus pairs with respect to body shape and colouration (Fig. 1). However, we refrained from analyses such as spectroradiometric assessments of body coloration (e.g., Dugatkin & Godin, 1996; Jordan et al., 2004) so as to avoid stressful anaesthesia or other forms of handling before the mate choice tests. Following this procedure, we successfully assigned 54 stimulus pairs (*n* = 27 per sex).

**Figure 1.**
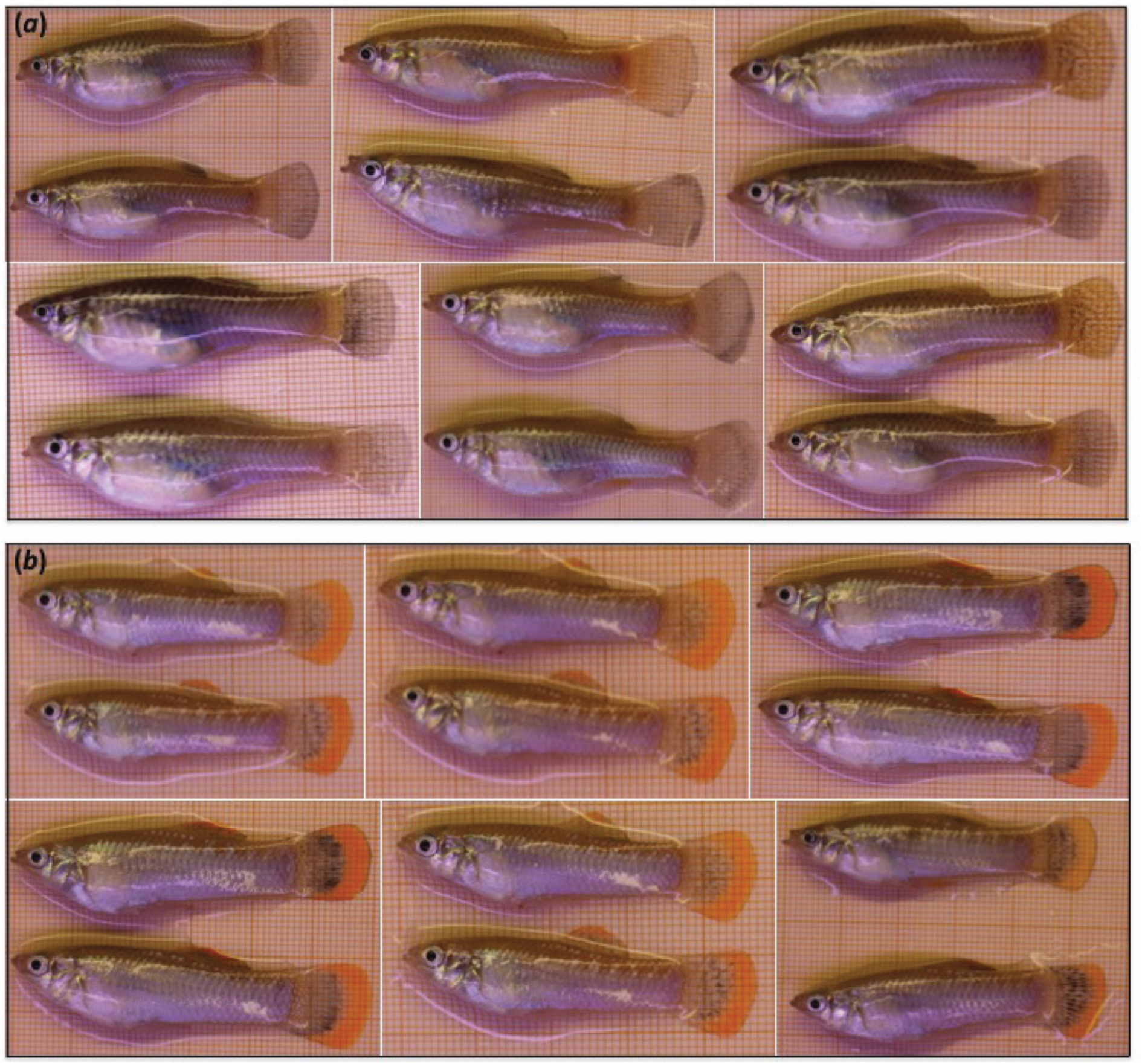
Examples of stimulus pairs (*a* females; *b* males), matched for standard length, body shape and colouration.

Of the 60 fish that were initially selected to serve as potential focal individuals, we randomly selected *n* = 27 individuals per sex as focal individuals for our mate choice tests and tested each focal individual with a different stimulus pair. To keep stress levels before the mate choice test as low as possible, SL of the focal fish was measured only after the mate choice tests. Upon completion of the behavioural tests, all fish were retransferred into their original stock tanks.

### Mate choice tests

During the association preference tests, a focal individual could choose between a risk-taking and a risk-averse mating partner. The test tank (60 × 25 × 35 cm) was divided into a central neutral zone (30 cm) and two lateral preference zones (15 cm each) adjacent to the stimulus compartments, which were separated from the main tank by transparent Plexiglas sheets (see Sommer-Trembo et al., 2016 for details). The focal fish was allowed to move freely between zones during a 5-min observation period, during which we scored times spent in either of the preference zones. We then switched side assignments of both stimulus individuals to avoid potential side biases and repeated measurement of association times. We summed times spent in association with either stimulus individual during the entire 10-min testing period.

We calculated strength of preference (SOP)-scores for risk-taking mating partners as: (time spent with risk-taking stimulus – time spent with risk-averse stimulus) / total association time. Thus, an SOP-score of +1 reflects maximal preference for the risk-taking stimulus and - 1 maximal preference for the risk-averse stimulus fish.

### Statistical analyses

All statistical tests were conducted using SPSS version 24.0. Where parametric tests were used, dependent variables met the assumptions of normal error distribution and homoscedasticity. Analyses were conducted separately for males and females.

To test for a directional preference for risk-taking mating partners, we used paired *t*-tests and compared association times near both types of stimulus individuals. We compared risk-taking scores and SOP-values between sexes using non-parametric Mann-Whitney *U*-tests. To test for potential effects of choosing individuals’ own personality (assortative mate choice), we ran univariate General Linear Models (GLM) using SOP-scores as the dependent variable and focal individuals’ risk-taking behaviour as a covariate. Due to limited sample sizes, we could not include all potentially biologically meaningful additional explanatory variables (size difference of risk-taking stimulus – risk-averse stimulus; focal individuals’ SL). However, when those covariates were included alongside focal individuals’ risk-taking behaviour in alternative GLMs, neither their main effects (*F*_1,24_ < 0.54, *P* > 0.47) nor interactions with risk-taking scores (*F*_1,23_ < 0.70, *P* > 0.41) were significant. Hence, in our main GLMs, the mean SL of the stimulus individuals and focal individuals’ risk-taking scores served as covariates. We excluded the non-significant interaction terms (females: *F*_1,23_ = 0.58, *P* = 0.46; males: *F*_1,23_ = 2.19, *P* = 0.15).

## Results

Males tended to be more risk-taking than females (Mean ± S.E. risk-taking-scores, females: 31.2 ± 4.6 s; males: 42.5 ± 3.4 s), but the difference was not statistically significant (Mann-Whitney *U*-test: *z* = 1.60, *p* = 0.11). Focal females showed a directional preference for risk-taking males and spent 263.4 ± 20.9 s in association with the risk-taking and 151.7 ± 16.9 s near the risk-averse stimulus male (paired *t*-test: *t*_26_ = 3.31, *p* = 0.003; Fig. 2*a*). By contrast, focal males did not show a directional preference related to females’ propensity to take risks (time spent with risk-taking female: 266.8 ± 23.0 s; with risk-averse female: 220.9 ± 22.8 s; paired *t*-test: *t*_26_ = 1.04, *p* = 0.31; Fig. 2*b*). However, strength of preference (SOP)-scores did not differ significantly between sexes (Mann-Whitney *U*-test: *z* = 1.54, *p* = 0.12).

**Figure 2.**
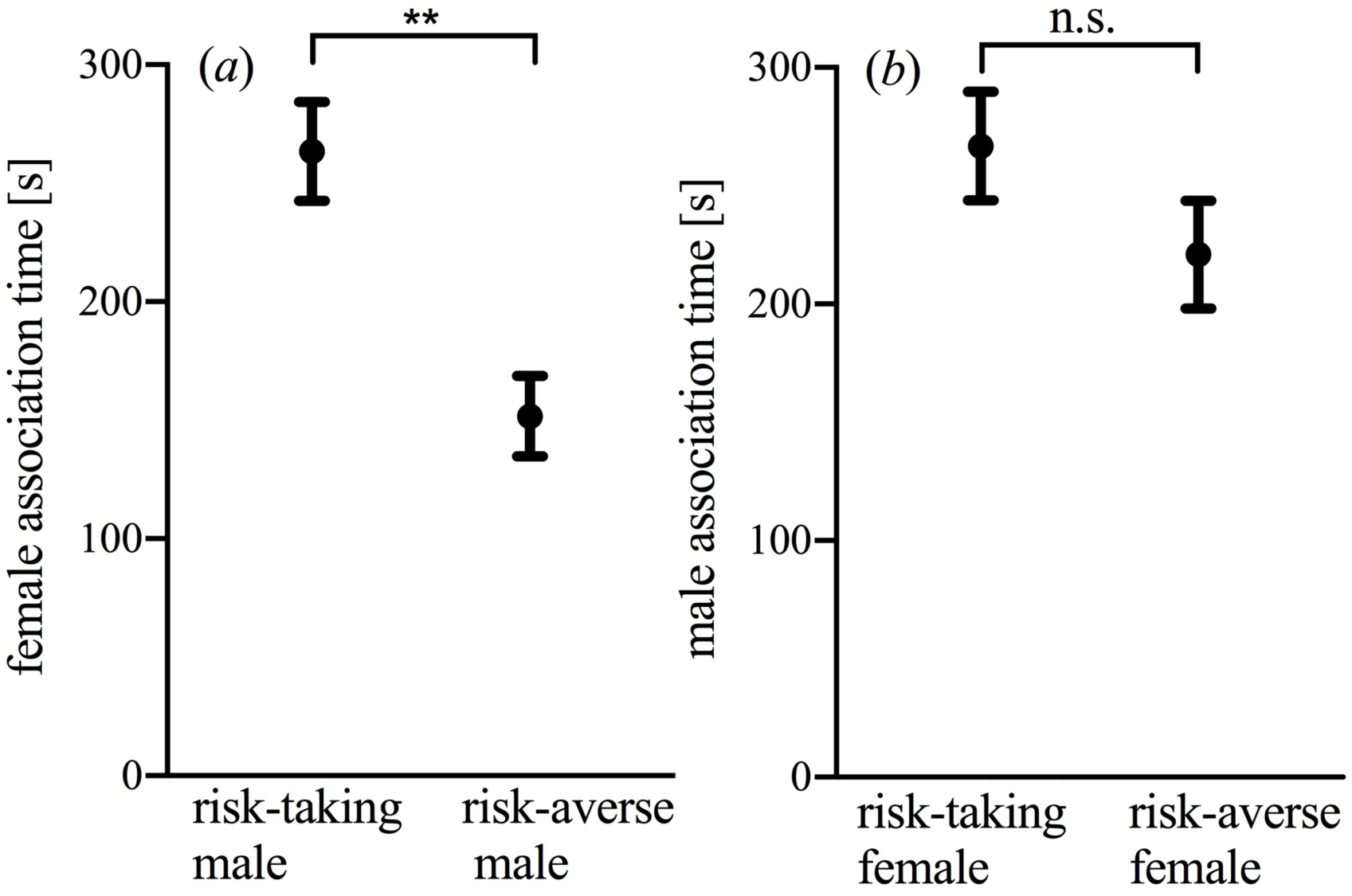
Results of dichotomous preference tests to assess directional mating preferences for risk-taking mating partners. Shown are the mean (± S.E.) times focal individuals (*a* females, *b* males) spent associating with risk-taking and risk-averse stimulus individuals of the opposite sex.

Focal females’ risk-taking tendency had a significant effect on their SOP (GLM, *F*_1,24_ = 4.94, *p* = 0.036; mean SL of stimulus males: *F*_1,24_ = 2.28, *p* = 0.14). A *post-hoc* Pearson correlation confirmed a significant, positive correlation between both variables (Fig. 3*a*). Neither focal males’ risk-taking tendency (*F*_1,24_ = 0.56, *p* = 0.46; Fig. 3*b*) nor the stimulus females’ mean body size (SL; *F*_1,24_ = 0.21, *p* = 0.65) had statistically significant effects on males’ SOP.

**Figure 3.**
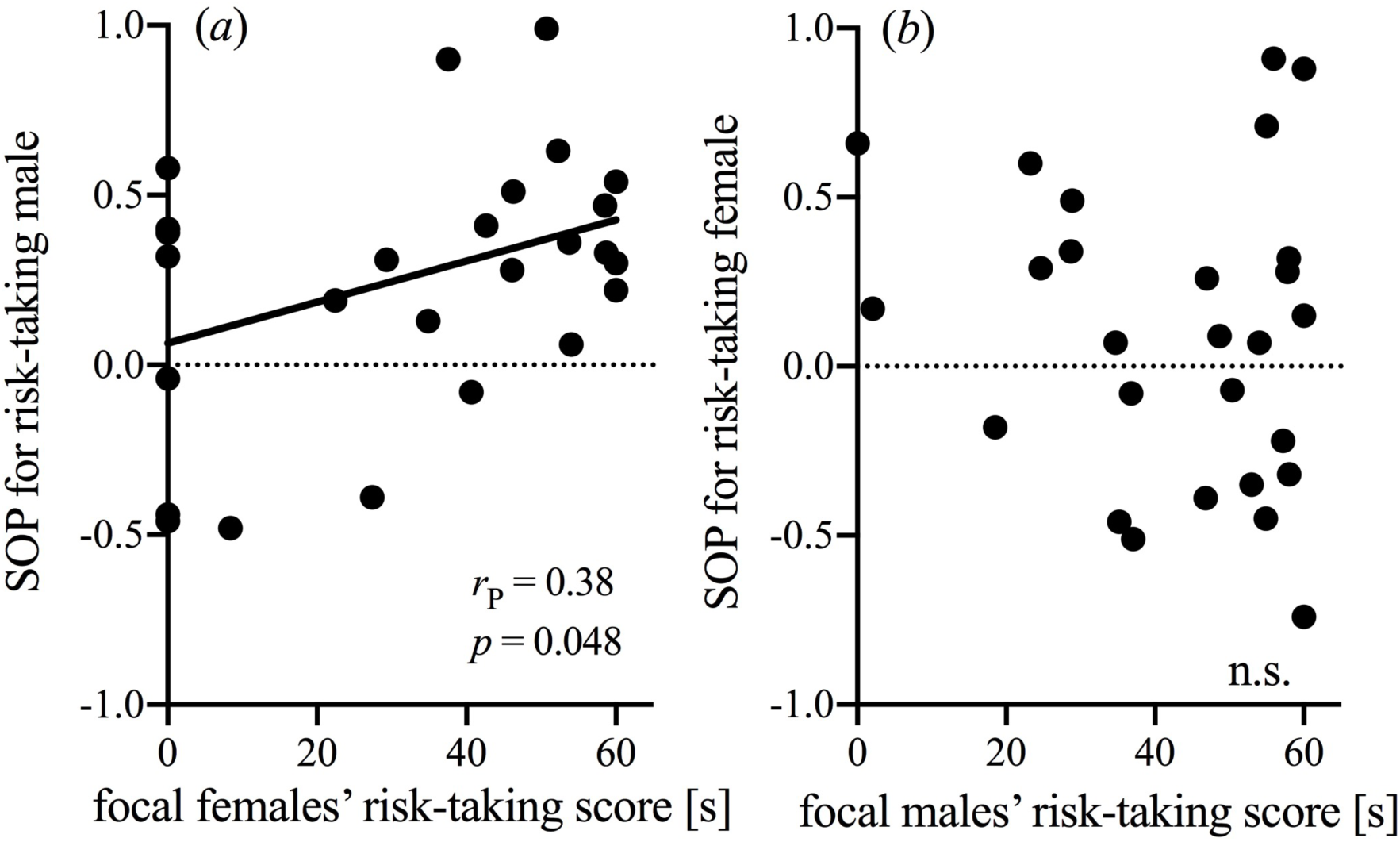
Scattergrams showing the correlation between focal individuals’ own risk-taking scores and their strength of preference (SOP) for risk-taking mating partners (testing for assortative mate choice). Females (*a*) but not males (*b*) showed a pattern where the choosing individual’s risk-taking tendency predicted variation in SOP-values, and risk-taking females showed stronger preferences for risk-taking stimulus males (*post-hoc* Pearson correlation).

## Discussion

Animal personality represents a major component of intraspecific phenotypic variation (Wolf & Weissing, 2012), but whether and how sexual selection (e.g., through mate choice) affects personality distributions remains understudied (Schuett, Tregenza & Dall, 2010). Using the livebearing fish *Poecilia mexicana*, we investigated whether a directional mating preference for risk-taking mating partners provides those individuals with a reproductive advantage (Godin & Dugatkin, 1996; Kortet, Niemelä, Vainikka & Laakso, 2019) and/or if the strength of preference for risk-taking individuals would be dependent on the choosing individuals’ own tendency to take risks (assortative mate choice; Scherer, Kuhnhardt & Schuett, 2017). We found a pattern in which both preference functions appear to interact: female (but not male) *P. mexicana* generally preferred risk-taking over risk-averse mating partners, but the strength of preference (SOP) for risk-taking males was dependent on the choosing females’ own personality type (i.e. risk-taking females exhibited stronger preferences for risk-taking males than risk-averse females).

A multitude of studies on mate choice considered mean preferences across individuals and inferred the existence of directional preferences for certain phenotypic traits (e.g., Kodric-Brown, 1985; Maan & Cummings, 2009; Marler & Ryan, 1997). In those cases, it often remains difficult to explain how additive genetic variance of the traits under sexual selection is maintained in natural populations (Brooks & Endler, 2001; Hoekstra et al., 2001; Morris, Nicoletto & Hesselman, 2003; Gasparini, Serena & Pilastro, 2013). Our results suggest that effects of the choosing individuals’ personality type could contribute to the maintenance of this variation, as they produce individual variation in mating preferences. Future studies in this and other species will need to consider the fact that the effects we describe here can easily be overlooked when research merely focusses on (more obvious) directional preferences, neglecting the potential drivers/correlates of individual variation in those preferences.

In our current study, personality differentially affected female and male mate choice. We argue that the adaptive significance of choosing risk-taking mating partners differs between sexes: in group-living animals, risk-taking is often associated with aggression and dominance (Colléter & Brown, 2011; Dahlbom, Lagman, Lundstedt-Enkel, Sundström & Winberg, 2011). Social dominance, in turn, can be a correlate of mating success in fish and other animals, especially in males (Ellis 1995; Jacob, Evanno, Renai, Sermier & Wedekind, 2009; Paull et al., 2010). In *P. mexicana*, dominant males monopolize and defend groups of females (Bierbach et al., 2014), and females receive less sexual harassment from those males (Plath, Parzefall & Schlupp, 2003). Choosing risk-taking mating partners, therefore, likely provides both direct and indirect (genetic) benefits to females. Moreover, females likely base their mate choice on certain phenotypic traits of males that are correlated with/indicative of risk-taking (including behaviour, see below), and the strength of this correlation could simply be weaker or absent in females.

Why did risk-taking females show a stronger preference for risk-taking males than risk-averse ones? One possible explanation would be that a trade-off between benefits of mating with risk-taking males and reproductive benefits of assortative mating (Ariyomo & Watt, 2013; Both, Dingemanse, Drent & Tinbergen, 2005; Kralj-Fišer, Sanguino Mostajo, Preik, Pekár & Schneider, 2013; Scherer, Kuhnhardt & Schuett, 2017) explains females’ mate choice. Moreover, risk-taking males tend to be more aggressive (Sih, Bell & Johnson, 2004) and risk-taking females could be more willing to accept the risk of interacting with aggressive males than risk-averse females.

Our results prompt the question of how exactly females discriminated between risk-taking and risk-averse males. We carefully matched stimulus males for morphological traits known to be involved in mate assessment. Nevertheless, females could differentiate between bold and shy males within the short time (10 min) of our behavioural tests. We may have overlooked subtle variation of certain (non-behavioural) traits that might correlate with differences in risk-taking tendencies, such as the intensity of male sexual ornamentation, but we consider this explanation unlikely. We argue in favour of another explanation: while females could not assess males’ responses to predators (Godin & Dugatkin 1996; Scherer, Kuhnhardt & Schuett 2017), variation in risk-taking likely correlates with other behavioural traits that females could evaluate during mate choice, especially males’ swimming patterns (Kern, Robinson, Gass, Godwin & Langerhans, 2016; Wilson & Godin, 2009), body posture, or time to emerge after disturbance (Brown, Jones & Braithwaite, 2005; Sommer-Trembo & Plath, 2018; this study), as slight disturbances occurred during the mate choice tests through handling, e.g. when switching stimulus males between the lateral compartments. Tracking programmes based on deep learning are currently being developed, which will enable us to analyse movement patterns of fish in unparalleled detail (e.g. Graving et al., 2019). It would be desirable to conduct a follow-up study using this state-of-the-art technology to investigate what components of movement patterns characterize different personality types and how these affect mate choice decisions.

Overall then, while mate choice based on directional preferences (Godin & Dugatkin 1996; Kortet, Niemelä, Vainikka & Laakso 2012; Reaney & Backwell) and assortative mate choice (Ariyomo & Watt, 2013; Both, Dingemanse, Drent & Tinbergen, 2005; Kralj-Fišer, Sanguino Mostajo, Preik, Pekár & Schneider, 2013; Scherer, Kuhnhardt & Schuett, 2017) seem to be mutually exclusive mechanisms, our results suggest that the existence of a directional preference does not preclude effects of choosing individuals’ personality on individual variation in mating preferences.

## Ethics

The experiments comply with the current laws on animal experimentation of the Federal Republic of Germany (*Regierungspräsidium Darmstadt* V-54-19c-20/15-F104/Anz.18).

## Data accessibility

The dataset supporting this article has been uploaded as part of the supplementary material.

## Authors’ contributions

C.S.-T. and M.S. collected data; C.S.-T. and M.P. conceived the idea for the analysis and analysed data; C.S.-T. and M.S. wrote the manuscript; all authors gave final approval for publication.

## Acknowledgements

We would like to thank Bruno Streit and Holger Geupel for their support to this study.

